# Chronic chemogenetic activation of hippocampal progenitors enhances adult neurogenesis and modulates anxiety-like behavior and fear extinction learning

**DOI:** 10.1101/2023.07.07.548060

**Authors:** Megha Maheshwari, Aastha Singla, Anoop Rawat, Toshali Banerjee, Sthitapranjya Pati, Sneha Shah, Sudipta Maiti, Vidita A. Vaidya

## Abstract

Adult hippocampal neurogenesis is a lifelong process that involves the integration of newborn neurons into the hippocampal network, and plays a role in cognitive function and the modulation of mood-related behavior. Here, we sought to address the impact of chemogenetic activation of adult hippocampal progenitors on distinct stages of progenitor development, including quiescent stem cell activation, progenitor turnover, differentiation and morphological maturation. We find that hM3Dq-DREADD-mediated activation of nestin-positive adult hippocampal progenitors recruits quiescent stem cells, enhances progenitor proliferation, increases doublecortin-positive newborn neuron number, accompanied by an acceleration of differentiation and morphological maturation, associated with increased dendritic complexity. Behavioral analysis indicated anxiolytic behavioral responses in transgenic mice subjected to chemogenetic activation of adult hippocampal progenitors at timepoints when newborn neurons are predicted to integrate into the mature hippocampal network. Furthermore, we noted an enhanced fear memory extinction on a contextual fear memory learning task in transgenic mice subjected to chemogenetic activation of adult hippocampal progenitors. Our findings indicate that hM3Dq-DREAD-mediated chemogenetic activation of adult hippocampal progenitors impacts distinct aspects of hippocampal neurogenesis, associated with the regulation of anxiety-like behavior and fear memory extinction.

## 1. Introduction

Adult hippocampal neurogenesis is an exquisitely regulated process, influenced by a variety of extrinsic and intrinsic factors (Zhao et al., 2008; Balu and Lucki, 2009; Bond et al., 2015; Toda et al., 2019). Extrinsic factors such as enriched environment, voluntary physical exercise, hippocampal-dependent learning tasks, chronic stress, dietary perturbations, as well as pharmacological agents such as antidepressants and anxiolytics are all known to influence adult hippocampal neurogenesis (Kempermann et al., 1997; Gould et al., 1999; Van Praag et al., 1999; Santarelli et al., 2003; Jiang et al., 2005; Stangl and Thuret, 2009; Snyder et al., 2011). The regulation of adult hippocampal neurogenesis could involve both direct effects mediated via the regulation of signaling pathways in adult hippocampal progenitor cells, as well as effects mediated via modulation of the neurogenic niche (Bond et al., 2015). The addition of newborn neurons within the adult hippocampal neurogenic niche throughout life has been implicated in diverse hippocampal functions, including the modulation of mood-related behavior, top-down control of stress-responsive neuroendocrine axes, as well as the regulation of learning and memory (Deng et al., 2010; Snyder et al., 2011; Anacker and Hen, 2017).

Several neurotransmitters have been implicated in the regulation of adult hippocampal neurogenesis, including but not restricted to the monoamines, glutamate, GABA, acetylcholine and distinct neuropeptides (Berg et al., 2013). Most of these neurotransmitters signal via the recruitment of G-protein-coupled receptors (GPCRs), several of which are reported to be expressed by adult hippocampal progenitors (Di Giorgi Gerevini et al., 2004; Kaneko et al., 2006; Giachino et al., 2014). While studies thus far have explored the neurogenic effects of select neurotransmitters and their specific GPCRs (Vaidya et al., 2007), there is limited evidence on the consequences of chemogenetic perturbation of Gq-, Gs-, or Gi-coupled signaling directly within adult hippocampal progenitors, and the associated effects on both adult hippocampal progenitor development and hippocampal-neurogenesis associated behaviors. A previous study showed that enhanced Gq signaling via chemogenetic activation of the hM3Dq-DREADD (Designer Receptors Exclusively Activated by Designer Drugs) in adult hippocampal progenitors evokes enhanced progenitor cell proliferation and a reduction in anxiety- and despair-like behavior, when assessed shortly after the chemogenetic activation paradigm (Tunc-Ozcan et al., 2019). This was the first evidence that chemogenetic perturbation of GPCR-linked signaling cascades within adult hippocampal progenitors can directly impact specific stages of adult hippocampal neurogenesis, and also influence neurogenesis-associated behaviors.

Adult hippocampal neurogenesis encompasses distinct stages of developmental progression, including quiescent stem cell activation and proliferation, followed by progenitor cell fate specification, maturation, migration, and functional integration into the hippocampal network (Kempermann et al., 2004, 2015; Ming and Song, 2005; Abbott and Nigussie, 2020). Radial glia-like stem cells, also known as Type-1 cells or RGLs, express markers such as nestin, sex determining region Y-box 2 (Sox2) and glial fibrillary acidic protein (GFAP) (Kempermann et al., 2004). These cells divide asymmetrically, giving rise to intermediate progenitor cells (IPCs, Type-2 cells), which are highly proliferative transit-amplifying cells and are categorized into: Type-2a cells, which express nestin and GFAP, and Type-2b cells, which express nestin, polysialylated-neural cell adhesion molecule (PSA-NCAM) and doublecortin (DCX). The Type-2 cells then transition into Type-3 cells, which are negative for nestin and positive for immature neuron markers PSA-NCAM and DCX. En route to forming mature granule cell neurons that functionally integrate into hippocampal dentate gyrus (DG) subfield network, newborn neurons transiently express the marker calretinin. The distinct stages of adult hippocampal progenitor development are characterized by specific markers that are used to evaluate various aspects of adult hippocampal neurogenesis (Hodge and Hevner, 2011; Von Bohlen Und Halbach, 2011; Beckervordersandforth et al., 2015; Zhang and Jiao, 2015).

In the present study, we have addressed whether enhancing hM3Dq-DREADD-mediated chemogenetic activation of Nestin-GFP-positive adult hippocampal progenitors alters their developmental progression, impacting proliferation, differentiation, and maturation of adult-born neurons. We find that hM3Dq-DREADD-mediated activation of nestin-positive adult hippocampal progenitors enhances progenitor turnover, accelerates differentiation, and morphological maturation, associated with a reduction in anxiety-like behavior and enhanced fear memory extinction, at time-points when the increased progenitor pool is predicted to functionally integrate into the hippocampal network and impact neurogenesis-associated behaviors.

## 2. Materials and Methods

### 2.1. Animals

Bigenic Nestin-rtTA::TetO-hM3Dq male mice were used for all experiments. The Nestin-rtTA transgenic mice (Yu et al., 2005), a kind gift from Prof. Steven Kernie (Department of Pediatrics, Columbia University, USA), were crossed to TetO-hM3Dq mice (Cat. No. 014093; Tg (TetO-CHRM3*)1Blr/J; Jackson Laboratories, USA), and bigenic genotypes were confirmed by PCR analysis. All animals were bred in the Tata Institute of Fundamental Research (TIFR, India) animal house facility, and maintained on a 12 h light/dark cycle from 7:00 A.M. to 7:00 P.M. with *ad libitum* access to food and water. Experimental procedures were conducted as per the guidelines of the Committee for the Purpose of Control and Supervision of Experiments on Animals, Government of India, and were approved by the TIFR animal ethics committees. Across all experiments, care was taken to minimize any pain or suffering and to restrict the number of animals used.

### 2.2. Drug treatments

To drive the expression of the hM3Dq-DREADD transgene in nestin-positive hippocampal progenitors, doxycycline (Dox, 1mM; Sigma−Aldrich, India) was administered in drinking water with 1 mM sucrose for four weeks to all Nestin-rtTA::TetO-hM3Dq bigenic mice, commencing at four weeks of age. Doxycycline administered Nestin-rtTA::TetO-hM3Dq bigenic mice received either vehicle (0.9% saline) or the DREADD ligand, clozapine-N-oxide (CNO, 2 mg/kg; Tocris, UK), intraperitoneally every alternate day for the duration of two weeks commencing at six weeks of age. To address the influence of CNO-mediated hM3Dq-DREADD activation of nestin-positive hippocampal progenitors on hippocampal progenitor proliferation, the mitotic marker 5-Bromo-2’-deoxyuridine (BrdU, 100 mg/kg; Sigma-Aldrich) was administered on the final day of CNO treatment, and mice were sacrificed two hours post-BrdU treatment. To examine the influence of CNO-mediated hM3Dq-DREADD activation on the maturation of hippocampal progenitors, vehicle and CNO-treated Nestin-rtTA::TetO-hM3Dq bigenic mice received three injections of BrdU (100 mg/kg) across three days, overlapping with the final three days of CNO/vehicle treatment and were sacrificed seven days post the final BrdU injection. Additional experimental cohorts of Nestin-rtTA::TetO-hM3Dq bigenic mice were subjected to vehicle or CNO treatment from six weeks of age for two weeks and subjected to behavioral analyses at three weeks after the cessation of CNO treatment.

### 2.3. Immunohistochemistry

Nestin-rtTA::TetO-hM3Dq bigenic mice were sacrificed by transcardial perfusion with 4% paraformaldehyde (PFA, Sigma-Aldrich). Coronal sections (40 μm) through the rostro-caudal extent of the hippocampus were generated using a vibratome (Leica VT1000 S, Germany), and processed for immunohistochemistry. For BrdU immunohistochemistry, every sixth section was processed for DNA denaturation and acid hydrolysis, followed by primary antibody rat anti-BrdU (1:200, Accurate Biochemicals, USA) incubation overnight at 4°C. Sections were then incubated with a secondary antibody (biotinylated goat anti-rat IgG, 1:500, Vector Laboratories, USA) for 2h followed by incubation with avidin-biotin complex (Vector Laboratories) for 1h, and visualized with diaminobenzidine (Sigma-Aldrich). For BrdU detection in immunofluorescence experiments, sections were incubated with Alexa 555-conjugated streptavidin (1:500, Invitrogen, Germany) for 1h after secondary antibody incubation.

We processed hippocampal sections to assess the numbers of transit-amplifying hippocampal progenitors immunopositive for the transcription factor T-box brain protein 2 (rabbit anti-Tbr2, 1:500, Abcam, USA), immature neurons immunopositive for doublecortin (DCX; rabbit anti-DCX; 1:250; Cell Signaling, USA) and adult hippocampal progenitors labeled with the nestin-GFP transgene (mouse anti-GFP; 1:500; Abcam). Sections were then incubated with appropriate secondary antibodies, Alexa Fluor 568-conjugated goat anti-rabbit (1:500; Invitrogen) and Alexa 488-conjugated donkey anti-mouse IgG (1:500, Invitrogen). The dendritic morphology of adult-born granule cell neurons was assessed using a primary antibody against PSA-NCAM (mouse anti-PSA-NCAM, 1:10,000; a kind gift form Prof. Tatsunori Seki, Juntendo University, Japan), followed by labeling with the secondary antibody, Alexa Fluor 488-conjugated donkey anti-mouse (1:500; Invitrogen).

For double immunofluorescence to assess the expression of the HA-tagged hM3Dq transgene in nestin-positive hippocampal progenitors (Nestin-GFP^+^-HA^+^), the transit-amplifying neural precursors (Nestin-GFP^+^-MCM2^+^) and immature neurons (BrdU^+^-Calretinin^+^), sections were incubated overnight with the following primary antibody cocktails: (1) goat anti-GFP (1:500; Abcam) and rat anti-HA (1:100; Roche, Switzerland) (2) mouse anti-GFP (1:500; Abcam) and rabbit anti-MCM2 (1:500; Abcam) and (3) rat anti-BrdU (1:200, Accurate Biochemicals) and rabbit anti-calretinin (1:250, Abcam, USA). Following washes, sections were incubated with secondary antibody cocktails: (1) Alexa Fluor 488-conjugated donkey anti-goat (1:500; Invitrogen) and Alexa Fluor 568-conjugated goat anti-rat (1:500; Invitrogen), (2) Alexa 488-conjugated donkey anti-mouse IgG (1:500, Invitrogen), and Cy5-conjugated goat anti-rabbit IgG (1:500, Invitrogen) and (3) biotinylated goat anti-rat IgG (1:500, Vector Labs) and anti-rabbit (1:500, Invitrogen). For triple immunofluorescence to assess the numbers of type 1 (Nestin-GFP^+^, GFAP^+^, DCX^-^), type 2a (Nestin-GFP^+^, GFAP^-^, DCX^-^) and type 2b (Nestin-GFP^+^, GFAP^-^, DCX^+^) hippocampal progenitors, sections were incubated with a cocktail of primary antibodies, goat anti-GFP, mouse anti-GFAP (1:500; Abcam), and rabbit anti-DCX (1:250; Cell Signaling). Sections were then incubated with a secondary antibody cocktail, Alexa Fluor 488-conjugated donkey anti-goat (1:500; Invitrogen), Alexa Fluor 568-conjugated goat anti-mouse (1:500; Invitrogen) and Cy5-conjugated goat anti-rabbit IgG (1:500, Invitrogen) for 2 hours. Sections were mounted using Vectashield (Vector Laboratories) and imaged using a Zeiss LSM 510 Exciter Laser Scanning microscope.

### 2.4. Cell counting analysis

Cell counting analysis was performed by an experimenter blind to the treatment conditions. The numbers of BrdU^+^, Tbr2^+^ and DCX^+^ cells within the subgranular zone (SGZ)/granule cell layer (GCL) of the DG subfield were estimated using a modified unbiased stereological approach. Cell counting analysis for each animal was performed in eight sections, 240 µm apart, across the rostral-caudal extent of the hippocampus as described previously (Malberg et al., 2000). The number of Nestin^+^ hippocampal progenitors per mm^2^ within the SGZ/GCL was determined across four to six sections spanning the rostrocaudal extent of the hippocampus. Neuronal morphology of PSA-NCAM positive cells was determined by neuronal tracing followed by Sholl analysis (NeuronStudio Beta). Percent colocalization of nestin-GFP with GFAP, DCX, or MCM2 markers was determined by analyzing 100 nestin-GFP-positive cells per animal (n = 4/group). Percent colocalization of BrdU with calretinin was determined by analyzing 50 BrdU-positive cells per animal (n = 4/group). Colocalization was confirmed using z-plane sectioning (1µm) on a Zeiss LSM 510 Exciter Laser Scanning microscope.

### 2.5. Cell culture and Calcium imaging

Dissociated hippocampal progenitor cell cultures were generated from hippocampi derived from mouse pups on postnatal day 1 (P1). Briefly, hippocampal progenitor cells were cultured in DMEM/F-12 with N2 supplement (Invitrogen) and 40 ng/ml FGF-2 (R&D Systems) on poly-lysine (Sigma) coated 35 mm glass bottom dishes (Nunc). On day four of plating, cells were subjected to calcium imaging using indo-1, AM (Invitrogen), a fluorescence probe undergoing a large shift in fluorescence emission maximum from 475 nm to 400 nm upon binding to Ca^2+^. We employed a confocal microscope (LSM 710, Carl Zeiss) coupled with a Ti:sapphire laser (MaiTai DeepSee, Spectra Physics) and custom modified for multiphoton imaging (Das et al., 2017) for the two-photon ratiometric measurement of calcium in cells before and after treatment with CNO. Cells were excited with a 740 nm high repetition rate (80 MHz) femtosecond pulsed (∼100 fs) laser, and the fluorescence signal was split into two channels—the transmitted channel (> 450 nm, signal from the Ca^2+^-free state of Indo-1: the “unbound channel”) and the reflected channel (< 450 nm, signal from the Ca^2+^-bound state of Indo-1: the “bound channel”). Cells were incubated with Hank’s balanced salt solution (Invitrogen) containing ratiometric calcium indicator Indo-1, AM (100 nM; Invitrogen) for 10 mins at 37°C, and then washed, prior to analysis of fluorescence in bound and unbound channels using the z-scan utility of the ZEN microscopy software (Carl Zeiss), in both the before CNO and after CNO (100 µM; 5 min; Tocris) conditions. The analysis was performed by selecting the soma of individual cells as regions of interest and then computing the average brightness of individual channels using Image J.

### 2.6. Behavioral tests

Independent experimental cohorts of Nestin-rtTA::TetO-hM3Dq bigenic mice were subjected to vehicle or CNO treatment from six weeks of age for a two-week duration, and were subjected to behavioral analyses at three weeks after the cessation of CNO treatment. The choice of this time-point was based on prior studies that indicate that the impact of newborn neurons on hippocampal neurogenesis-associated behaviors requires the functional integration of these newborn neurons into mature hippocampal networks, and this has been estimated to be at time-points of three to six weeks of adult hippocampal progenitor development (Kempermann et al., 2004; Toni and Schinder, 2015). Vehicle and CNO-treated Nestin-rtTA::TetO-hM3Dq bigenic mice were subjected to open field test (OFT) and elevated plus maze (EPM) to assess anxiety-like behavior, and to the fear conditioning paradigm to assess fear memory. For the OFT, mice were released into one corner of the open field arena (40 cm x 40 cm x 40 cm), chosen at random, and were allowed to explore for five minutes. The behavior was recorded using an overhead camera and automated behavioral tracking analysis was performed using Ethovision 3.1 (Noldus, Netherlands). The total distance traveled, percent distance traveled in the center, percent time spent in the center, and number of entries to the center of the arena were calculated. The EPM was a plus shaped platform with opposing open and closed arms (45 cm x 10 cm each) elevated 50 cm above the ground. Mice were introduced into the arena facing the open arms, and behavior was recorded for five mins using an overhead camera. Behavioral tracking was performed using Ethovision 3.1 (Noldus, Netherlands) and the total distance traveled, percent distance traveled in the open arms, and percent time spent in the open arms, was determined. The fear conditioning paradigm was conducted in chambers with internal dimensions of 25 cm x 25 cm x 25 cm (Panlab Harvard Apparatus, USA). A house light and fan mounted directly above the chamber provided illumination and white noise, respectively. Freezing was defined as the complete absence of motion for a minimum of 0.5 s. The fear conditioning procedure was conducted over three days. On day zero, mice were placed in the conditioning chamber and baseline freezing was assessed. On day one, mice were placed in the conditioning chamber and received pairings between a shock (2s, 0.3mA) and a tone (80 dB) in the training context ‘A’ (square walls, textured floor, vanilla scent. Each mouse was placed into the chamber for three minutes and shock was administered once. Freezing was scored throughout the three-minute-long training. On day two, to test fear recall, mice were introduced in training context ‘A’ for three minutes, without presenting any shock and freezing was scored. To test the extinction of fear memory, mice were exposed to training context ‘A’ without a shock, for 8 days, for a duration of three minutes every day and the entire session was scored for freezing.

### 2.7. Statistical analysis

Results were subjected to statistical analysis using the program Prism 5 (GraphPad, San Diego, CA, USA). Experiments with two groups were analyzed for differences using the unpaired Student’s *t* test, with significance determined at *p* < 0.05. Experiments that measured progression across time were subjected to repeated measures two-way analysis of variance (ANOVA) analysis followed by the Bonferroni *post-hoc* test. Significance was determined at values of *p* < 0.05.

## 3. Results

### 3.1. Selective expression of the HA-tagged hM3Dq-DREADD in nestin-positive adult hippocampal progenitors

We established the bigenic Nestin-rtTA::TetO-hM3Dq mouse line to address the influence of hM3Dq-DREADD mediated chemogenetic activation of hippocampal progenitors on diverse aspects of hippocampal neurogenesis, including neuronal proliferation, survival, and maturation, as well as addressing the influence on anxiety-like behavior. Capitalizing on a reverse-tTA-based strategy, we administered the tetracycline analog, doxycycline, to bigenic Nestin-rtTA::TetO-hM3Dq mice (Fig. 1A), to drive the expression of the HA-tagged hM3Dq- DREADD transgene in nestin-positive adult hippocampal progenitors (Fig. 1B). Double immunofluorescence studies and confocal microscopic analyses for the HA-tag and nestin-GFP, revealed colocalization of the HA-tagged hM3Dq-DREADD in nestin-GFP- immunopositive adult hippocampal progenitors within the SGZ region of the DG subfield (Fig. 1C). We did not observe the presence of the HA-tagged hM3Dq-DREADD transgene in cells that lacked nestin-GFP expression within the hippocampal neurogenic niche. We next addressed the influence of the exogenous DREADD ligand, CNO on calcium levels in dispersed hippocampal progenitor cultures, immunopositive for the HA-tagged hM3Dq-DREADD (Fig. 1D). Dispersed hippocampal progenitor cultures were incubated with the ratiometric intracellular calcium indicator, Indo-1, AM, followed by calcium imaging prior to CNO application and immediately following CNO exposure (Fig. 1E). Quantitative analysis indicated that the ratio of bound to unbound calcium was significantly enhanced post-CNO treatment in hippocampal progenitors derived from Nestin-rtTA::TetO-hM3Dq bigenic mice (Fig. 1F). Taken together, these *in vitro* and *in vivo* results indicate the selective expression of the HA-tagged hM3Dq-DREADD in nestin-GFP-positive adult hippocampal progenitors in Nestin-rtTA::TetO-hM3Dq bigenic mice, and indicate that upon CNO-mediated hM3Dq-DREADD activation, hippocampal progenitors exhibit an increase in intracellular calcium levels.

**Figure 1.**
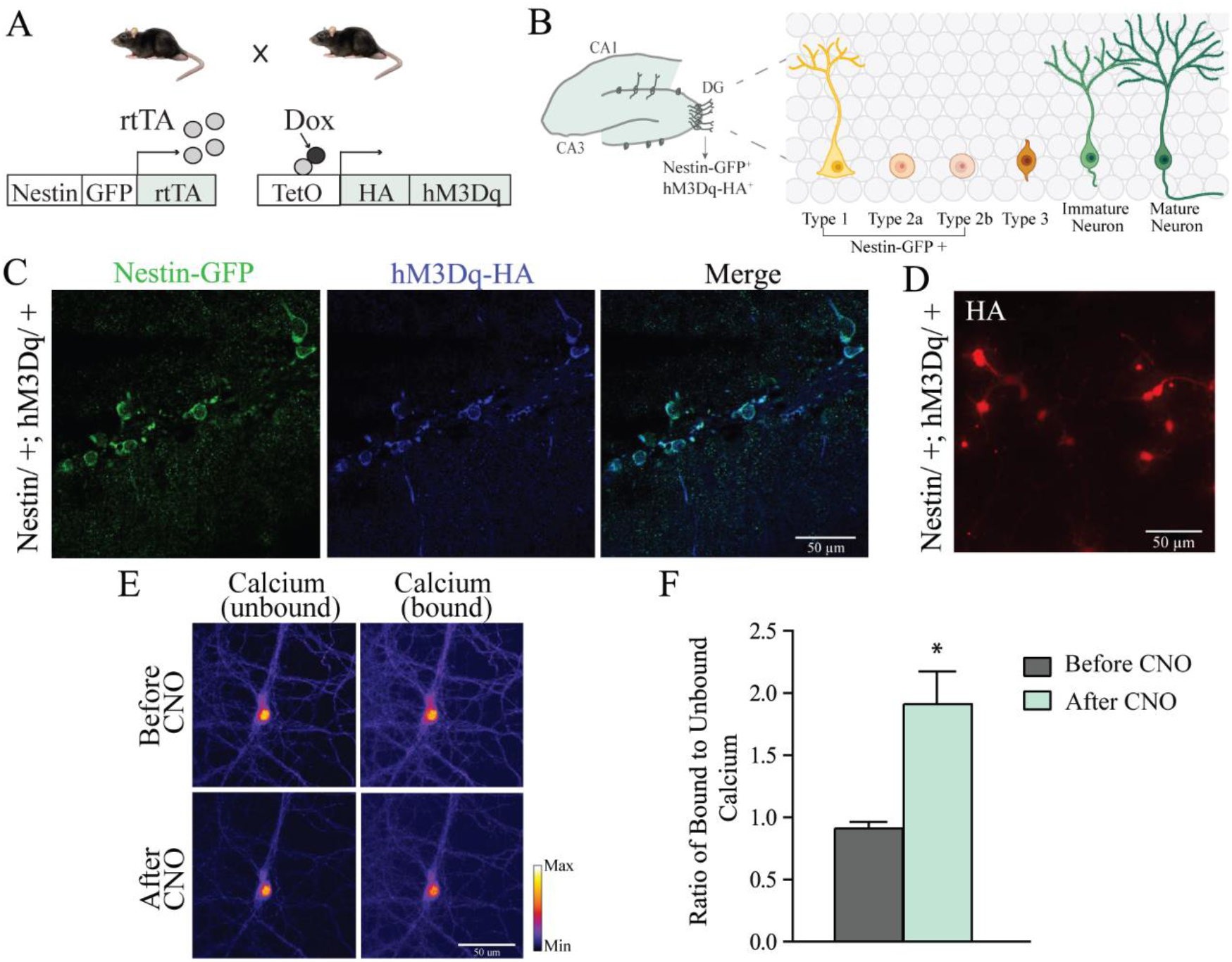
Selective expression of the HA-tagged hM3Dq-DREADD in nestin-positive adult hippocampal progenitors. A, Shown is a schematic of the experimental strategy for the generation of Nestin-rtTA::TetO-hM3Dq bigenic mice to selectively drive the expression of the hM3Dq-DREADD in nestin-GFP-positive adult hippocampal progenitors. rtTA – reverse tetracycline transactivator. Doxycycline was administered to the bigenic mice from the age of 4 to 6 weeks to drive the HA-tagged-hM3Dq transgene expression in nestin-positive progenitors. B, Shown is a schematic of the dentate gyrus (DG) hippocampal subfield indicating the stage-specific progression of markers associated with adult hippocampal progenitor development, with Type 1 progenitors that are nestin and GFAP double-immunopositive, Type 2a progenitors that are nestin-positive, and Type 2b progenitors that are nestin and DCX double-immunopositive. The HA-tagged hM3Dq-DREADD transgene is expressed within nestin-immunopositive adult hippocampal progenitors. C, Shown are representative confocal images indicating the expression of the HA-tagged hM3Dq-DREADD colocalizing with the nestin-GFP expression in adult hippocampal progenitors within the DG subfield (n = 4). D, Shown is a representative image indicating the expression of the HA-tagged hM3Dq-DREADD in dispersed hippocampal progenitor cells derived from Nestin-rtTA::TetO-hM3Dq bigenic mouse pups at postnatal day 1. E, Shown are representative images of dispersed hippocampal progenitor cells treated with the Ca^2+^-sensitive fluorophore Indo-1, AM, with fluorescence intensity ratios (bound/unbound) indicative of a change in unbound and bound calcium, both before and after treatment with the DREADD ligand, CNO. F, Shown is the quantitation of the ratio of bound to unbound calcium in hippocampal progenitor cells before and after treatment with the DREADD ligand CNO. Results are expressed as mean values ± SEM, n = 16 neurons/treatment condition compiled across N = 3, \**p* < 0.05 as compared to control (Paired Student’s *t*-test). Scale bar, 50 μm.

### 3.2. Chronic hM3Dq-DREADD mediated activation of nestin-positive hippocampal progenitors enhances adult hippocampal progenitor proliferation and stage-specific progenitor marker expression

We subjected Nestin-rtTA::TetO-hM3Dq bigenic mice, maintained on doxycycline from four weeks of age to induce expression of the hM3Dq-DREADD transgene in nestin-positive hippocampal progenitors, to treatment with the DREADD ligand CNO commencing from six weeks of age for a period of two weeks (Fig. 2A). To assess the effects of hM3Dq-DREADD mediated activation of nestin-positive hippocampal progenitors on progenitor turnover, we treated both CNO and vehicle administered Nestin-rtTA::TetO-hM3Dq bigenic mice with the mitotic marker BrdU (100 mg/kg), and sacrificed them two hours later (Fig. 2A). Quantitative analysis revealed a significant increase in the number of BrdU-positive progenitors within the DG subfield in the CNO-treated Nestin-rtTA::TetO-hM3Dq bigenic mice as compared to their vehicle-treated controls (Fig. 2B, C).

**Figure 2.**
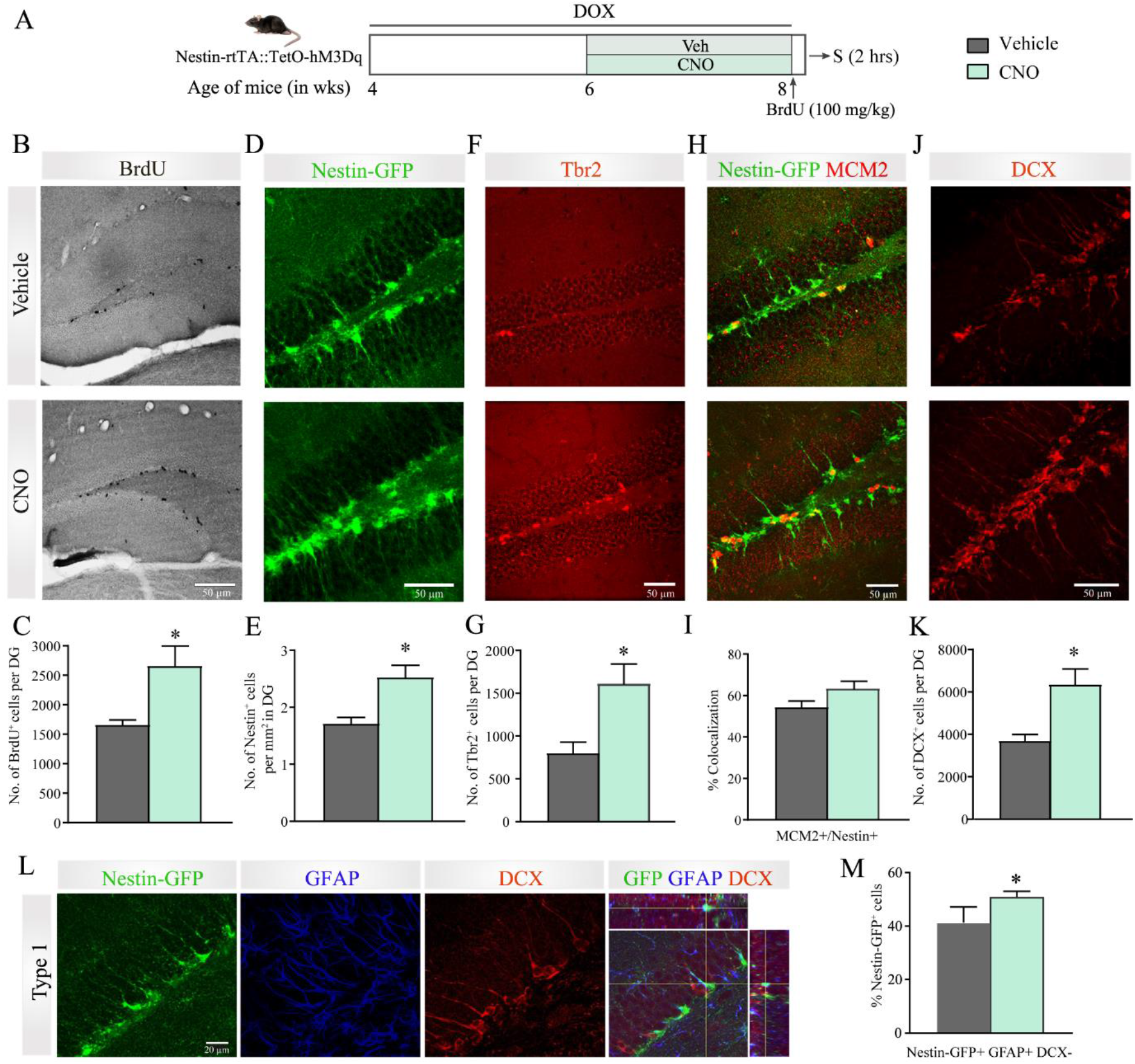
Chronic hM3Dq-DREADD mediated activation of nestin-positive hippocampal progenitors enhances adult hippocampal progenitor proliferation and stage-specific progenitor marker expression. A, Shown is a schematic of the experimental paradigm where Nestin-rtTA::TetO-hM3Dq bigenic mice were maintained on doxycycline, commencing at 4 weeks of age to induce the hM3Dq-DREADD transgene in nestin-positive progenitors, were subjected to treatment with the DREADD agonist, CNO (2 mg/kg) or vehicle for a duration of 2 weeks starting from 6 weeks of age. Nestin-rtTA::TetO-hM3Dq bigenic mice were administered the mitotic marker BrdU (100 mg/kg) at the end of CNO/vehicle treatment and sacrificed 2 hours later. B, Shown are representative brightfield images from the vehicle and CNO-treated Nestin-rtTA::TetO-hM3Dq bigenic mice for BrdU-positive adult hippocampal progenitor cells within the dentate gyrus (DG) subfield. C, Quantitative analysis revealed a significant increase in BrdU-positive cell number in the DG of CNO-treated Nestin- rtTA::TetO-hM3Dq mice as compared to the vehicle group. Shown are representative confocal images of Nestin-GFP positive (D), Tbr2-positive (F), Nestin-GFP-MCM2 double-positive cells (H), and DCX-positive cells (J) from the vehicle and CNO-treated Nestin-rtTA::TetO- hM3Dq bigenic mice. Scale bars: 50 µm. Quantitative analysis revealed a significant increase in Nestin-GFP-positive (E), Tbr2-positive (G), and DCX-positive (K) cells in the DG of CNO- treated mice as compared to controls. The percent colocalization of nestin with MCM2 (I) was not found to be altered between the CNO-treated Nestin-rtTA::TetO-hM3Dq bigenic mice, as compared to the vehicle-treated controls. L, Shown are representative confocal images indicating triple-immunofluorescence staining for Nestin-GFP (green), GFAP (blue), and DCX (red) to distinguish Type 1 (GFP^+^GFAP^+^DCX^-^), Type 2a (GFP^+^GFAP^-^DCX^-^) and Type 2b (GFP^+^GFAP^-^DCX^+^) progenitor cells in the DG subfield. Scale bars: 20 µm. M, Quantification of Type 1, Type 2a, and Type 2b cells as percentage of Nestin-GFP positive progenitors revealed a significant increase in Type 1 progenitors in the CNO-treated mice as compared to vehicle-treated controls. All results are expressed as the mean ± SEM (n = 5 per group). \**p* <0.05 compared to controls (unpaired Student’s *t*-test).

We next assessed whether chronic hM3Dq-DREADD mediated activation of nestin-positive adult hippocampal progenitors alters the numbers of the nestin-positive progenitor pool, and determined the numbers of nestin-positive cells/mm^2^ in the hippocampal neurogenic niche (Fig. 2D). A significant increase in the numbers of nestin-positive cells/mm^2^ was noted in CNO-treated Nestin-rtTA::TetO-hM3Dq bigenic mice (Fig. 2E). Following this, we proceeded to assess the influence of hM3Dq-DREADD mediated activation of nestin-positive hippocampal progenitors on the expression of stage-specific markers for adult hippocampal progenitors within the neurogenic niche. The impact of chemogenetic activation of adult hippocampal progenitors on the total numbers of Tbr2-positive hippocampal progenitors, which label the intermediate-stage progenitors (Hodge et al., 2012), considered to be the transit-amplifying cells in the hippocampal neurogenic niche, was determined. We noted a significant increase in the numbers of Tbr2-positive hippocampal progenitors within the DG subfield of CNO-treated Nestin-rtTA::TetO-hM3Dq bigenic mice as compared to their controls (Fig. 2F, G). To examine the consequences of chronic chemogenetic activation of nestin-positive hippocampal progenitors on the number of DCX-immunopositive progenitors and immature neurons within the hippocampal neurogenic niche, we performed DCX immunohistochemistry (Fig. 2J). Cell counting analysis indicated a significant increase in the number of DCX-positive cells within the DG subfield in Nestin-rtTA::TetO-hM3Dq bigenic mice subjected to chronic treatment with the DREADD ligand, CNO as compared to bigenic mice treated with vehicle (Fig. 2K). We then addressed whether the enhanced numbers of nestin-positive progenitors noted following chemogenetic activation of the nestin-pool of progenitors, is also reflective of enhanced exit from quiescent stages by examining colocalization of nestin with MCM2, which labels progenitors that have exited quiescence (Zhang and Jiao, 2015) (Fig. 2H). Confocal analysis for the percent colocalization of nestin with the MCM2 marker did not reveal any differences between vehicle and CNO-treated Nestin-rtTA::TetO-hM3Dq bigenic mice (Fig. 2I).

We next sought to address the impact of chemogenetic activation of nestin-positive adult hippocampal progenitors on the fraction of Type 1 (Nestin-GFP and GFAP double-positive) progenitor cells in the total progenitor pool (Fig. 2L). Triple immunofluorescence studies and confocal-based colocalization analysis revealed a significant increase in the Nestin- GFP and GFAP double-positive Type 1 hippocampal progenitors in Nestin-rtTA::TetO- hM3Dq bigenic mice subjected to chronic treatment with CNO as compared to vehicle-treated controls (Fig. 2M), suggesting an increase in the quiescent neural progenitor (QNP) pool. This result suggests that the enhanced numbers of nestin-positive progenitors noted following chronic chemogenetic activation of the nestin-positive progenitor pool may likely reflect an increase in Type-1 adult hippocampal progenitors.

Collectively, these findings indicate that chronic hM3Dq-DREADD mediated activation of nestin-positive adult hippocampal progenitors evokes increased numbers of Type 1 progenitor cells, enhanced progenitor cell turnover as revealed by increased mitotic marker labeling, increased numbers on Tbr2-positive transit amplifying progenitors, as well as an increase in DCX-positive immature neurons.

### 3.3. Chronic hM3Dq-DREADD mediated activation of nestin-positive adult hippocampal progenitors enhances dendritic complexity and accelerates progenitor maturation

Given that we noted robust effects of chronic chemogenetic activation of nestin-positive progenitors on diverse aspects of adult hippocampal neurogenesis, we next sought to address whether this chemogenetic activation of adult hippocampal progenitors also influences the dendritic complexity of immature neurons and alters progenitor maturation. Nestin-rtTA::TetO-hM3Dq bigenic mice maintained on doxycycline from four weeks of age, received treatment with the DREADD ligand, CNO or vehicle for two weeks with the administration of BrdU overlapping with the final three days of CNO/vehicle treatment and were sacrificed a week later (Fig. 3A). Dendritic complexity of adult-born immature neurons, was assessed using immunofluorescence for PSA-NCAM, a marker for immature neurons within the hippocampal neurogenic niche (Zhang and Jiao, 2015), followed by neuronal tracing and Sholl analysis (Fig. 3B). Sholl analysis of PSA-NCAM-positive immature neurons revealed that chronic chemogenetic activation of nestin-positive adult hippocampal progenitors results in a significant increase in the number of intersections per µm distance from the soma (20 µm to 100 µm, *p* < 0.05) (Fig. 3C), the total number of intersections (Fig. 3D), and the total dendritic length (Fig. 3E), indicative of enhanced dendritic complexity in PSA-NCAM-positive adult-born immature neurons.

**Figure 3.**
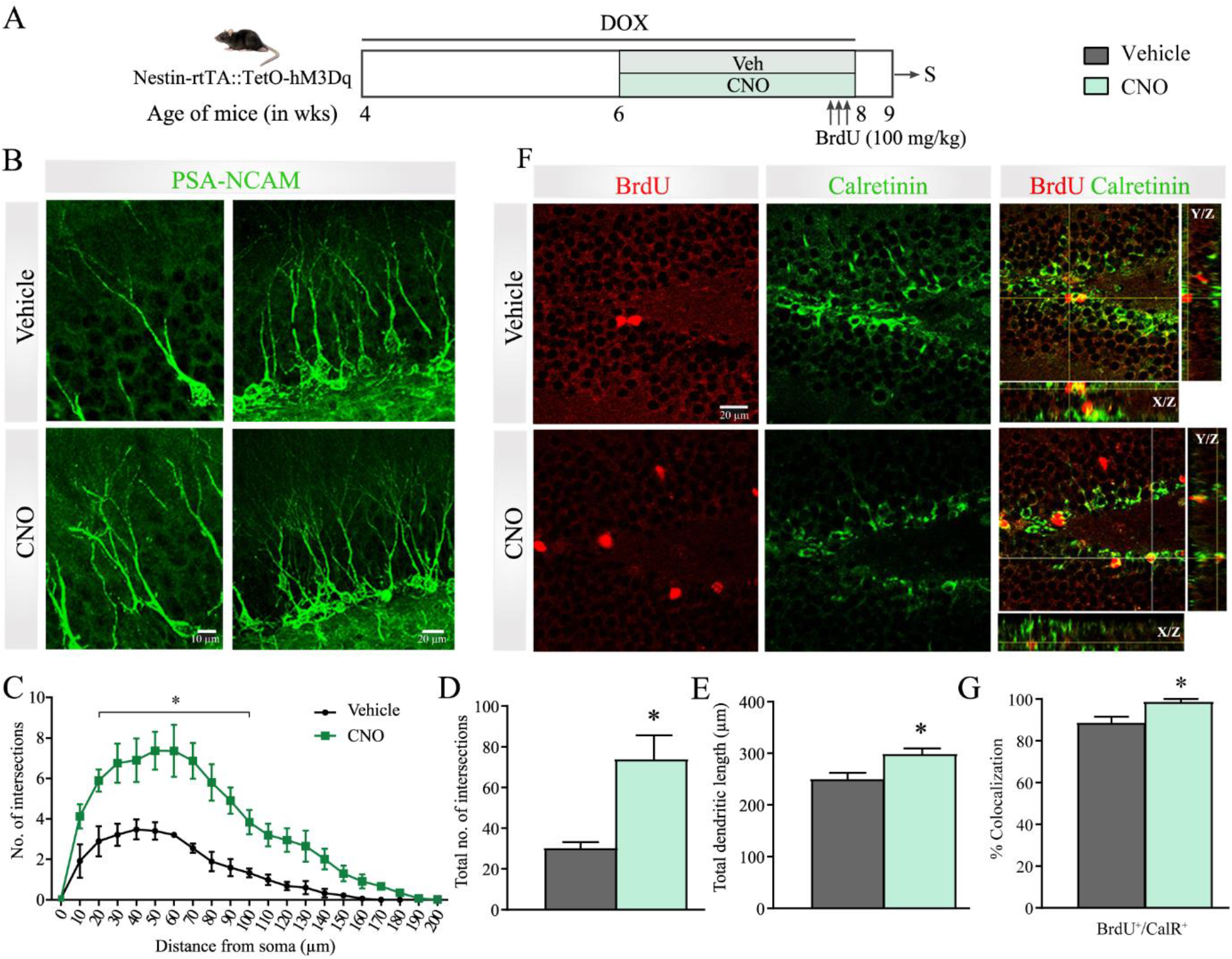
Chronic hM3Dq-DREADD mediated activation of nestin-positive adult hippocampal progenitors enhances dendritic complexity and accelerates progenitor maturation. A, Shown is a schematic of the experimental paradigm wherein Nestin-rtTA::TetO-hM3Dq bigenic mice maintained on doxycycline, commencing at 4 weeks of age were subjected to treatment with the DREADD agonist CNO or vehicle for 2 weeks starting from 6 weeks of age. Post vehicle or CNO treatment, mice received the mitotic marker BrdU once daily for three days and were sacrificed 7 days post-BrdU treatment. B, Shown are representative confocal images of PSA-NCAM labeled immature neurons in the DG subfield, indicating the dendritic arbors of immature neurons from the vehicle and CNO-treated Nestin-rtTA::TetO-hM3Dq bigenic mice. Scale bars: 10 μm (high magnification); 20 μm (low magnification). Shown is a quantification of the number of intersections per μm of distance from the soma (C), the total number of intersections (D), and the total dendritic length (E) across the dendritic arbor of PSA-NCAM stained neurons in the DG of vehicle and CNO-treated mice. (C) Sholl Analysis indicated that the number of intersections was significantly enhanced in the CNO-treated group at a distance of 20 - 100 μm from the soma (C). \**p* < 0.05 (Two-way Repeated Measures ANOVA, Bonferroni post-hoc multiple comparisons). The total number of intersections was significantly enhanced in the CNO-treated cohort as compared to the vehicle-treated control group (D) \**p* <0.05 (unpaired Student’s *t*-test), n = 15 neurons/animal, n = 3 mice/group. E, Total dendritic length was significantly increased in Nestin-rtTA::TetO-hM3Dq bigenic mice that received CNO treatment. \**p* < 0.05 (unpaired Student’s *t*-test), n = 15 neurons/animal, n = 3 mice per treatment group. F, Shown are representative confocal images indicating co-localization of BrdU with Calretinin (CalR), a transient marker expressed during adult hippocampal progenitor maturation. Scale bar: 20μm. G, Quantitative analysis revealed a significant increase in the percent colocalization of BrdU with CalR, in CNO-treated Nestin-rtTA::TetO-hM3Dq bigenic mice as compared to their vehicle-treated controls. Results are expressed as the mean ± SEM (n = 3-4 per group) \**p* < 0.05 (unpaired Student’s *t*-test).

We next examined whether the chronic chemogenetic activation of adult hippocampal progenitors influences their maturation. We sought to address whether the BrdU pulse-labeled adult hippocampal progenitors in CNO-administered Nestin-rtTA::TetO-hM3Dq bigenic mice exhibit a change in the degree of colocalization with calretinin, a marker transiently expressed in the early postmitotic steps of neuronal differentiation (Brandt et al., 2003), as compared to their vehicle-treated controls (Fig. 3F). We observed a small but significant increase in the percentage of BrdU and calretinin double-positive cells in the CNO-treated cohort as compared to the control group (Fig. 3G). Taken together, these findings suggest that chemogenetic activation of adult hippocampal progenitors enhances the dendritic complexity of immature neurons and may also accelerate their acquisition to stage-specific markers.

### 3.4. Chronic hM3Dq-DREADD mediated activation of nestin-positive hippocampal progenitors reduces anxiety-like behavior

Given we observed robust effects of chronic chemogenetic activation of nestin-positive progenitors on progenitor turnover, numbers of quiescent and transit amplifying progenitors, immature neuron number, maturation and dendritic complexity, we sought to address whether the enhanced hippocampal neurogenesis noted following chronic chemogenetic activation was associated with an influence on anxiety-like behavior. We chose to assess the behavioral consequences at the time-point of three weeks post the cessation of CNO-treatment, which was also the five-week time-point post commencement of CNO administration (Fig. 4A, F). The choice of this time-point was based on prior evidence that adult-born neurons that are three to four weeks of age, appear to effectively integrate and maximally influence the excitability of the DG network (Toni and Schinder, 2015). Further, adult-born dentate gyrus cells that are four to six weeks of age have been shown to influence behavioral tasks linked to anxiety-like behaviors and fear conditioning (Denny et al., 2012). Hence, we chose to examine the influence of chronic chemogenetic activation of nestin-positive progenitors on anxiety-like behaviors in the OFT (Fig. 4A-E) and EPM (Fig. 4F-J), at the three-week time point post cessation of a two week long, chronic CNO treatment.

**Figure 4.**
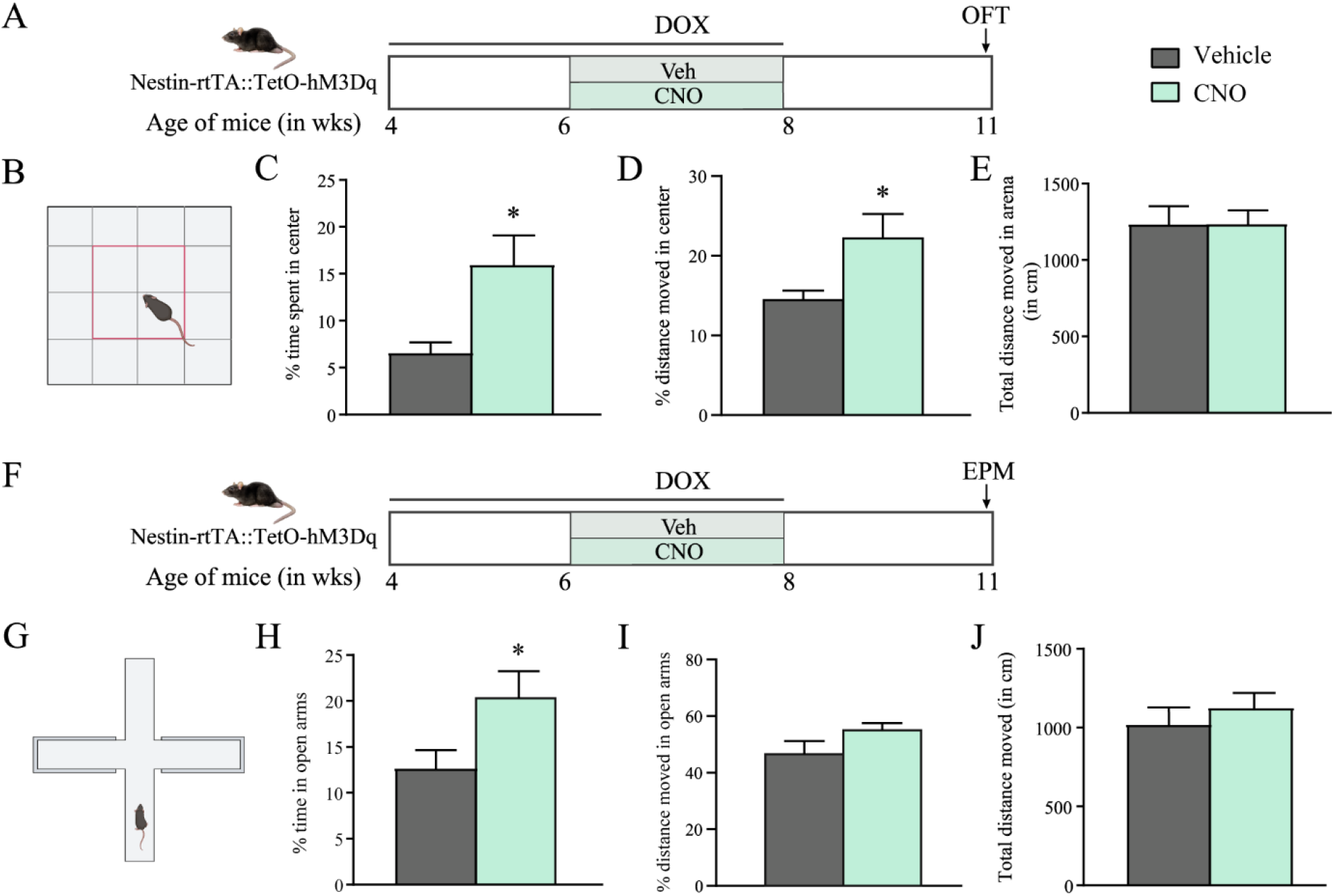
Chronic hM3Dq-DREADD mediated activation of nestin-positive hippocampal progenitors reduces anxiety-like behavior. Shown are schematics of the experimental paradigms followed to chemogenetically activate hM3Dq-DREADD in nestin-positive hippocampal progenitors using bigenic Nestin-rtTA::TetO-hM3Dq mice that were maintained on doxycycline from the age of 4 weeks to 8 weeks to switch on the hM3Dq-DREADD transgene, and then administered CNO or vehicle to chemogenetically activate the hM3Dq-DREADD from the age of 6 weeks to 8 weeks prior to being subjected to behavioral tests at 11 weeks, namely the open field test (OFT) (A) and elevated plus maze (EPM) test (F) in separate cohorts. (A - E) Mice were subjected to the OFT to assess anxiety-like behavior, and CNO-treated mice exhibited reduced anxiety-like behavior as noted by a significant increase in percent time (C) and percent distance (D) moved in the center of the OFT arena. Total distance moved in the arena was not altered (E). \**p* < 0.05 (unpaired Student’s *t*-test), n = 12-15 mice/ group. (F - J) Mice were subjected to the EPM task to assess anxiety-like behavior, and CNO-treated mice showed reduced anxiety-like responses indicated by a significant increase in the percent time (H). No significant change was observed in either percent distance (I) in the open arms and the total distance moved in the EPM arena (J). \**p* < 0.05 (unpaired Student’s *t*-test), n = 11-13 mice/ group. Results are expressed as mean ± SEM.

On the OFT (Fig. 4B), we noted a significant decline in anxiety-like behavior as revealed by an increase in percent time spent in the center (Fig. 4C), and percent distance moved in the center of the arena (Fig. 4D). We observed no differences in total distance moved in the arena (Fig. 4E). In independent cohorts, we also noted a significant decline in anxiety-like behavior on the EPM (Fig. 4G), as indicated by an increase in percent time spent in the open arms (Fig. 4H). We noted no difference in the percent distance moved in open arms (Fig. 4I) and total distance moved in the EPM (Fig. 4J), between vehicle and CNO-treated Nestin-rtTA::TetO-hM3Dq bigenic mice.

### 3.5. Chronic hM3Dq-DREADD activation of nestin-positive hippocampal progenitors in adulthood accelerates the extinction of fear memory

We next sought to address the influence of chemogenetic activation of adult hippocampal progenitors on contextual fear conditioning (CFC) behavior (Fig. 5A, B). We observed no difference in the baseline freezing (Fig. 4C) between the vehicle and CNO-treated Nestin-rtTA::TetO-hM3Dq bigenic mice, prior to exposure to the CFC paradigm. Further, the acquisition of conditioned fear was not altered across the two groups, as the percent of context-elicited freezing was comparable between the two treatment groups (Fig. 4D). This revealed that the ability to form a fear memory is not altered following chronic chemogenetic activation of adult hippocampal progenitors. We next exposed the vehicle and CNO-treated Nestin-rtTA::TetO-hM3Dq bigenic mice to fear extinction training, and noted facilitation of fear memory extinction in the CNO-treated cohort (Fig. 4E), as revealed by a significant reduction in percent freezing on days three to five of the extinction training paradigm.

**Figure 5.**
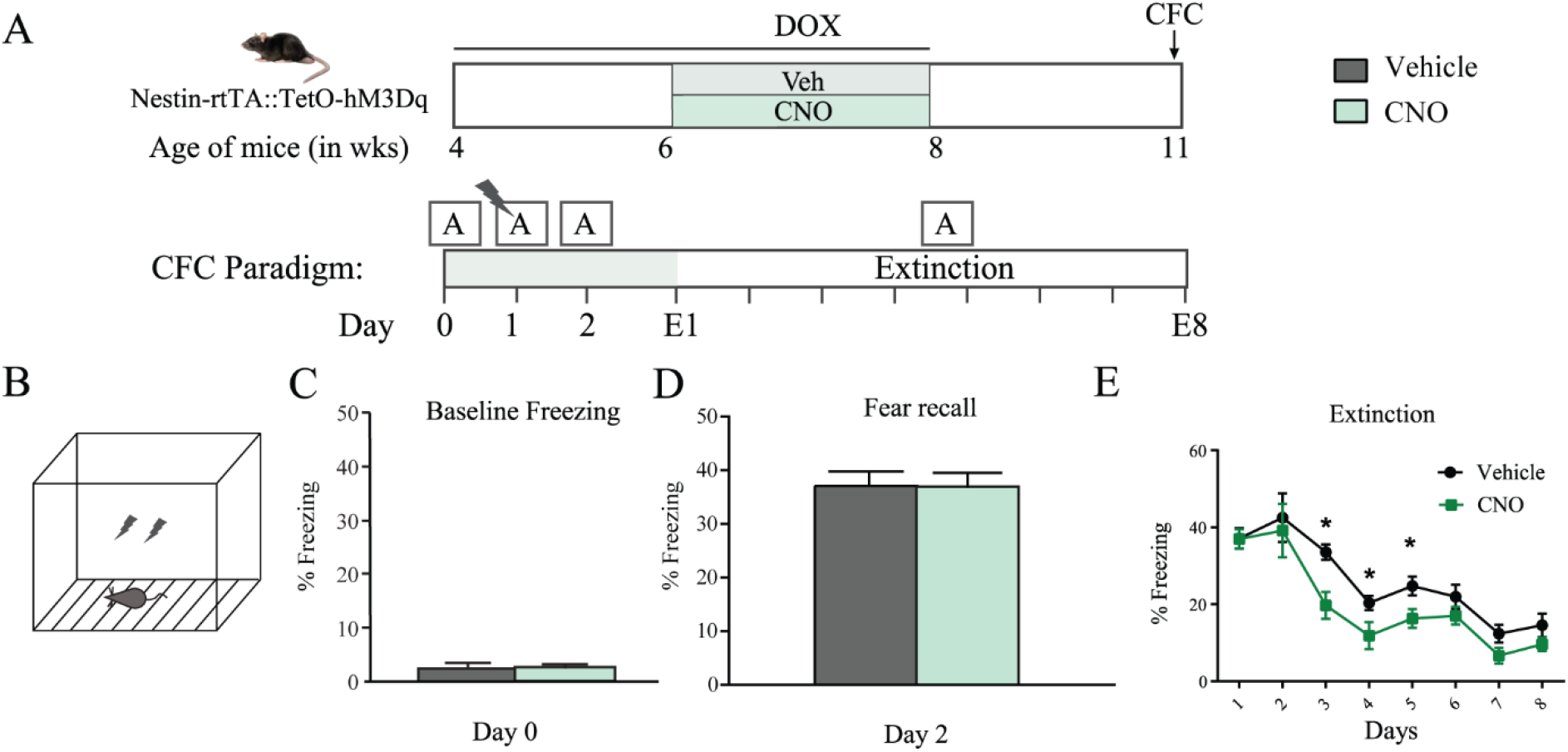
Chronic hM3Dq-DREADD activation of nestin-positive hippocampal progenitors in adulthood enhances fear memory extinction. A, Shown is a schematic of the experimental paradigm followed to chemogenetically activate the hM3Dq-DREADD in nestin-positive hippocampal progenitors using bigenic Nestin-rtTA::TetO-hM3Dq mice that were maintained on doxycycline from the age of 4 weeks to 8 weeks to switch on the hM3Dq-DREADD transgene, and then administered CNO or vehicle to chemogenetically activate the hM3Dq-DREADD from the age of 6 weeks to 8 weeks prior to being subjected to contextual fear conditioning (CFC) at 11 weeks. Mice were subjected to contextual fear conditioning, to assess fear memory acquisition and extinction. Analysis of baseline freezing on day 0 in context ‘A’, indicated no differences between the treatment groups (A, C). Animals were subjected to fear conditioning on day 1, wherein a shock paired with a tone was delivered to mice in context ‘A’ (intensity 0.3 mA, 2s, paired 5 times with a tone of 80 dB) (A, B). On day 2, no differences were noted in fear recall as revealed by a similar extent of freezing (D) exhibited by both vehicle and CNO treated groups on exposure to context ‘A’. Animals were then subjected to extinction training (A), wherein the CNO-treated group exhibited enhanced fear extinction, as indicated by reduced freezing behavior across day 3, 4, 5 of extinction training (E). \**p* < 0.05 compared to controls, two-way ANOVA with repeated measures, Bonferroni post-hoc multiple comparisons test; n = 10-12 mice/group.

## 4. Discussion

We find that hM3Dq-DREADD-mediated activation of adult hippocampal neuronal progenitors within the SGZ of the DG subfield influences distinct steps of the developmental progression of progenitors, and modulates hippocampal neurogenesis-responsive behavioral tasks (Revest et al., 2009; Seo et al., 2015). Our findings provide evidence that a chemogenetic approach to modulate Gq-coupled signaling in nestin-positive adult hippocampal progenitors can enhance quiescent stem cell activation, enhancing the pool of Type-1 hippocampal progenitors and promote progenitor cell division. Furthermore, hM3Dq-DREADD-mediated chemogenetic activation of nestin-positive hippocampal progenitors is associated with an acceleration of the acquisition of immature neuronal markers like calretinin, as well as an increased dendritic complexity of immature neurons. The neurogenic consequences of chemogenetic activation of adult hippocampal progenitors are correlated with a reduction in anxiety-like behaviors on ethologically relevant behavioral tasks, and with an increased fear memory extinction in a contextual fear learning paradigm, at time points when newborn neurons are predicted to integrate into mature hippocampal networks (Denny et al., 2012).

A prior study reported that hM3Dq-DREADD-mediated activation of Ascl1-positive hippocampal progenitors, that predominantly includes transit-amplifying neural progenitors (Type 2 cells), with the possibility of also recruiting a small subset of neural stem cells (Type 1 cells), leads to an increase in hippocampal progenitor proliferation and a reduction in anxiety-like behavior, immediately post the cessation of a three-week duration chemogenetic activation paradigm (Tunc-Ozcan et al., 2019). In contrast, the specific progenitor stages targeted in our study using the Nestin promoter to drive the hM3Dq-DREADD involves robust expression in both Type 1 and Type 2 cells. Nevertheless, our findings are in keeping with this prior report (Tunc-Ozcan et al., 2019) indicating that CNO-mediated activation of hM3Dq-DREADD increases progenitor proliferation, as revealed by enhanced numbers of dividing progenitors, labeled either with an endogenous proliferation marker such as Ki-67 used in the prior study, or with exogenous mitotic markers like BrdU, as used in our study. Previous reports suggest a link between diverse Gq-coupled GPCRs and the regulation of adult hippocampal neurogenesis (Vaidya et al., 2007), implicating the metabotropic glutamate receptor (Nochi et al., 2012), the muscarinic acetylcholine receptor M1 (Itou et al., 2011) and the 5-HT_2C_ receptor in the regulation of hippocampal progenitor proliferation (Banasr et al., 2004). Furthermore, calcium signaling pathways that lie downstream of Gq-coupled signaling are also implicated in the regulation of adult hippocampal progenitor turnover (Li et al., 2012; Toth et al., 2016; Xu et al., 2018). Our ratiometric calcium imaging analysis *in vitro* using hippocampal progenitor cultures derived from Nestin-rtTA::TetO-hM3Dq bigenic mouse pups indicates that CNO-mediated hM3Dq-DREADD activation recruits calcium signaling cascades reported to lie downstream of Gq-coupled signaling in progenitors. Our findings of enhanced progenitor proliferation in response to chemogenetic activation of hippocampal progenitors suggest that exogenous DREADD-based activation of Gq signaling in adult hippocampal progenitors phenocopies the observations from pharmacological studies targeting Gq-coupled GPCRs reported to be expressed in the hippocampal neurogenic niche (Itou et al., 2011; Nochi et al., 2012; Giachino et al., 2014)

In our study we have extensively characterized the impact of CNO-mediated hM3Dq- DREADD activation of nestin-positive hippocampal progenitors on distinct stages of the developmental progression of progenitors. We find that chemogenetic activation of nestin-positive adult hippocampal progenitors enhances the total number of BrdU-positive cells, reflective of increased hippocampal progenitor turnover. The mitotic marker BrdU would label all proliferating adult hippocampal progenitors, including the rapid amplifying Type 2 progenitor cells, but would also include the smaller fraction of proliferative neural stem cells (Type 1) within the hippocampal neurogenic niche. We find that chemogenetic activation of adult hippocampal progenitors evoked significantly higher numbers of both total Nestin-GFP- positive progenitors, which encompass multiple early stages (Type 1 and Type 2) of adult hippocampal progenitor development, as well as Tbr-2 positive progenitors, that label intermediate neuronal progenitors, predominantly Type 2 cells in the hippocampal neurogenic niche (Hodge et al., 2012). Using double immunofluorescence for MCM2 and Nestin, we assessed whether the proliferative pool (MCM2+) of Nestin-positive progenitors, predominantly the Type 2 cells, was enhanced as a fraction of the total number of Nestin-positive cells, and noted no difference. However, we observed a significant increase in the fraction of Nestin-positive cells that were GFAP-positive, but DCX-negative, which is a pattern of marker expression associated with quiescent stem cells (Type 1) in the neurogenic niche. Collectively, this suggests that upon chemogenetic activation of nestin-positive progenitors, distinct early stages of hippocampal progenitor development are impacted including progenitor cell division, as well as quiescent stem cell activation. A caveat of our study is that we are limited in our ability to parcellate the effects of chemogenetic activation on individual stages of the early developmental progression of hippocampal progenitors, given that our genetic driver nestin is expressed in several of these developmental stages. A recent intriguing study suggests that the balance between quiescent versus activated neural stem cells in adult neurogenic niches is governed by differences in calcium dynamics, with an increase in cytosolic calcium from IP3-mediated gating of intracellular stores regulating the process of stem cell activation (Gengatharan et al., 2021).

Commensurate with the enhanced proliferation of adult hippocampal progenitors noted upon CNO-mediated DREADD activation, we found a robust increase in the number of DCX- positive immature neurons. Furthermore, we noted an enhanced dendritic length, as well as increased dendritic complexity, of immature neurons within the hippocampal neurogenic niche, using PSA-NCAM as a marker to assess morphological complexity (Seki and Arai, 1993). A prior study addressing the impact of chemogenetic activation of newborn hippocampal neurons in a genetically modified Tau mouse model, indicated that in addition to rescuing the impaired dendritic arbors of newborn granule cells in mutant mice, chemogenetic activation was also capable of enhancing the dendritic complexity of newborn neurons in wild-type mice (Terreros-Roncal et al., 2019). Despite the differences in methodology, retroviral strategies versus a bigenic mouse model approach to target adult hippocampal progenitors, and variation in CNO treatment duration and the experimental paradigm, the commonalities of observation suggest that chemogenetic activation of adult hippocampal progenitors enhances dendritic complexity and morphological maturation of newborn neurons. The enhanced dendritic complexity raised the intriguing possibility of accelerated differentiation of newborn neurons following chronic hM3Dq-mediated activation of progenitors, this notion was supported by evidence of enhanced colocalization of BrdU with calretinin, a transient marker (Brandt et al., 2003) observed in the early stages of immature newborn neuron development in the adult hippocampal neurogenic niche. It has been suggested that accelerated maturation of immature neurons could also impact membrane excitability and shape the sculpting of both afferents to and efferents from adult-born granule cells (Piatti et al., 2006; Trinchero et al., 2021). Our findings raise the speculative possibility that chemogenetic activation of nestin-positive adult hippocampal progenitors could impact integration of these newborn neurons into the mature dentate gyrus network.

In our study we also examined the behavioral consequences of chemogenetic activation of adult hippocampal progenitors, choosing a time-point three weeks after the cessation of vehicle or CNO administration. Since our CNO treatment was itself for a duration of two weeks, the pool of newborn neurons exposed to CNO treatment would vary between three to five weeks of age. The selection of this paradigm was based on several previous studies which suggest that the critical period for maturation and integration of adult-born granule cells is between the second and sixth week of age of these newborn neurons (Denny et al., 2012; Toni and Schinder, 2015). During this temporal window newborn neurons exhibit enhanced excitability and synaptic plasticity, accompanied by a lowered threshold for long term potentiation strongly impacting the DG network (Aimone et al., 2010; David et al., 2010; Deng et al., 2010; Anacker and Hen, 2017; Huckleberry and Shansky, 2021; Vyleta and Snyder, 2023). We observed a significant anxiolytic effect on the OFT and EPM following chemogenetic activation of nestin-positive hippocampal progenitors. These findings are in agreement with a previous study that reported anxiolytic effects on the OFT following chemogenetic of Ascl1-positive adult hippocampal progenitors, albeit at an earlier time point of two hours following two weeks of CNO administration (Tunc-Ozcan et al., 2019). Several previous studies that have used both pharmacological and genetic perturbation strategies to enhance ongoing adult hippocampal neurogenesis and reported a reduction in anxiety-like behaviors (Bergami et al., 2009; Revest et al., 2009; Campos et al., 2013; Quesseveur et al., 2013; Kobayashi et al., 2014; Mohammad et al., 2018). However, there are also studies that suggest that simply increasing adult hippocampal neurogenesis using a genetic strategy to prevent newborn neuron death does not per se impact anxio-depressive behaviors, but can prevent the enhanced anxio-depressive behaviors evoked by chronic stress or corticosterone administration (Hill et al., 2015; Culig et al., 2017). In this regard our study differs, as we find that a chemogenetic approach that activates nestin-positive progenitors and enhances adult hippocampal neurogenesis can decrease anxiety-like behavior in the absence of a stressor paradigm. In keeping with prior evidence, our findings suggest that the decrease in anxiety-like behavior noted maybe a consequence of enhanced neurogenesis, with an increased number of newborn neurons integrating into the DG network (Dias et al., 2012; Hill et al., 2015; Mohammad et al., 2018; Li et al., 2022).

We also examined the consequences of chemogenetic activation of adult hippocampal progenitors on fear learning and extinction in the contextual fear conditioning paradigm (CFC), which has been previously shown to be sensitive to perturbations in the extent of ongoing adult hippocampal neurogenesis (Drew and Huckleberry, 2017). Prior studies indicate that a decline in the available pool of newborn granule cells results in both impaired acquisition of fear memory and extinction learning (Deng et al., 2009). Furthermore, previous studies suggest that four to six-week-old newborn neurons contribute to the acquisition of fear learning in the CFC paradigm (Denny et al., 2012). Our experimental paradigm commenced with the CFC protocol three weeks after the cessation of the CNO treatment, a time-point when the cohort of chemogenetically activated newborn neurons would be within the range of three to five weeks of age. While vehicle and CNO-treated mice showed comparable levels of fear learning in the CFC paradigm, we noted that CNO-treated mice with a chemogenetic activation of adult hippocampal progenitors exhibited a significant facilitation of fear extinction learning. Although the role of adult hippocampal neurogenesis in fear extinction is not fully understood, working models posit that two to four-week-old newborn neurons, which are inherently more excitable, appear to be critical in fear memory extinction, with a key role in contributing to ‘the discrimination between the original fear memory trace and the new safety memory originating during fear extinction’ (Agis-Balboa and Fischer, 2014). Our data that chemogenetic activation of adult hippocampal progenitors, which drives enhanced progenitor turnover, differentiation and morphological maturation, results in enhanced fear extinction learning at time-points when newborn neurons are within three to five weeks of age is in keeping with working models of the role of adult neurogenesis in fear extinction learning (Jaako-Movits et al., 2005; Ko et al., 2009; Pan et al., 2012; Fitzsimons et al., 2013; Kheirbek et al., 2013; Denny et al., 2014; Seo et al., 2015; Drew and Huckleberry, 2017; Huckleberry et al., 2018; Inoue et al., 2023).

In conclusion, we find that chemogenetic activation of nestin-positive adult hippocampal progenitors enhances progenitor cell division, increases quiescent stem cell activation, accelerates the acquisition of transient markers of immature neuron development and enhances the dendritic complexity of newborn neurons. Furthermore, in temporal windows when chemogenetically activated, newborn neurons are three to five weeks of age, we find a decline in anxiety-like behaviors and an enhancement of fear extinction learning. This study adds to the growing body of evidence that links modulation of newborn neuron formation in the hippocampal neurogenic niche with the regulation of mood-related behavior.

## CRediT authorship contribution statement

**Megha Maheshwari:** Visualization, Investigation. **Aastha Singla:** Investigation, Writing – original draft. **Anoop Rawat:** Investigation. **Sthitapranjya Pati:** Investigation. **Toshali Banerjee:** Investigation. **Sneha Shah:** Investigation. **Sudipta Maiti:** Investigation. **Vidita A. Vaidya:** Conceptualization, Writing – original draft, Writing – review & editing, Funding acquisition.

## Acknowledgments

This work was supported by intramural funding from the Tata Institute of Fundamental Research and the Department of Atomic Energy, Mumbai (RTI4003 to VV). We thank Dr. Shital Suryavanshi, K.V. Boby, and the animal house staff at the Tata Institute of Fundamental Research, Mumbai, for technical assistance.

## Author contributions

MM, AS, AR, SP, TB, SS contributed to animal experiments and data analysis. AS and VV wrote the manuscript.

